# APP overexpression delays synaptic development and alters neuronal network properties

**DOI:** 10.1101/2025.03.12.641184

**Authors:** Fulin Ma, Himanshu Akolkar, Ryad Benosman, Karl Herrup

## Abstract

The amyloid precursor protein (APP) regulates neuronal excitability by altering the structure of the axon initial segment. Using high density multielectrode arrays we compared the electrophysiological consequences of APP overexpression with exposure to β-amyloid (Aβ) and found that the two manipulations affected neural activity in largely non-overlapping ways. In mature cultures (>18 days in vitro, DIV18), APP reduced the firing probability of individual neurons, while β-amyloid stripped synapses impacting whole-network connectivity. In immature cultures (DIV10), increasing APP – either acutely with lentiviral vectors or genetically by culturing APP transgenic neurons – revealed a separate developmental function. Synaptogenesis and axonal branch formation were blocked. Aβ, by contrast, interfered primarily with network behavior. Our findings suggest that Alzheimer’s disease is likely an amalgamation of APP mis-regulation plus the effects of Aβ accumulation. They also urge consideration of the idea that familial and sporadic forms of Alzheimer’s may represent distinct disease processes.

## Introduction

The amyloid precursor protein (APP) is a multifunctional Type 1 transmembrane protein expressed in virtually all cell types including neurons. While APP has been intensively studied for its role as the parent of the β-amyloid (Aβ) peptide, an identifying feature of Alzheimer’s disease^1^, it also has important physiological roles in many neuronal functions. APP is detected as early as the pre-implantation stage^2^ and expression continues throughout development. It has been suggested to play a suppressive role in neurogenesis^3^ and is critical for proper neuronal migration^4-6^. APP also plays a role in axonal growth and guidance^7-10^, in dendritic maturation^11-13^ and in synaptogenesis^14-17^. These diverse roles have largely been identified in experiments where APP levels were altered acutely. By contrast, chronic elimination of APP (in *App*-null mutant mice) results in only mild phenotypes: cortical development is normal, and axonal connectivity is only slightly disrupted^18-21^. Apart from its developmental functions, APP plays a homeostatic role in tuning neuronal activity. It mediates this action through the regulation of the axon initial segment (AIS). The AIS is a specialized region of the axon where a high density of voltage gated ion channels favors action potential initiation^22-24^. Neuronal activity shifts the position and length of the AIS in such a way as to reduce action potential firing probability^24-26^. Unexpectedly, this activity-dependent modulation of neuronal activity requires APP^27^. Neuronal activity increases APP and increased APP, on its own, is sufficient to shorten the AIS and cause it to shift away from the cell body, with the expected decrease in neuronal activity confirmed by GCaMP. These data validate the effect of APP at the single cell level but leave unanswered several questions about how such changes might affect a larger neuronal network.

In the current work we have approached this question using high density multielectrode arrays (HD-MEA). Unlike traditional MEAs, HD-MEAs have inter-electrode distances of less than 20 µm, allowing for very precise detection of AIS location, action potential propagation and axon branching data. Using the high spatial and temporal resolution of neuronal activity afforded by these arrays, we have analyzed cultures of dissociated mouse embryonic cortical neurons and describe the properties of their networks as they develop. We then show that elevating the levels of APP delays network development differently depending on the timing of the manipulation, and that Aβ disrupts physiological activity in a distinct way, in keeping with its lack of impact on the length and location of the AIS^27^. The findings offer new insights into APP function and have important implications for our understanding of the biological basis of Alzheimer’s disease.

## Results

### The electrical properties of wild type neuronal cultures develop on independent timelines

Primary cortical neurons were isolated from C57BL/6J embryo cortices at 16.5 days of gestation (E16.5) and cultured on high density multi-electrode arrays (MEA) as described in the Methods (Fig. 1A). By 18 days in vitro (DIV18), the neurons had become electrically active, grown processes of considerable length, and established active synapses. Using MAP2 to reveal the location of the neurons relative to the electrodes (Fig. 1B), we found that a typical neuronal cell body was nearly equal in size to a single electrode and could lie in contact with up to four.

**Figure 1.**
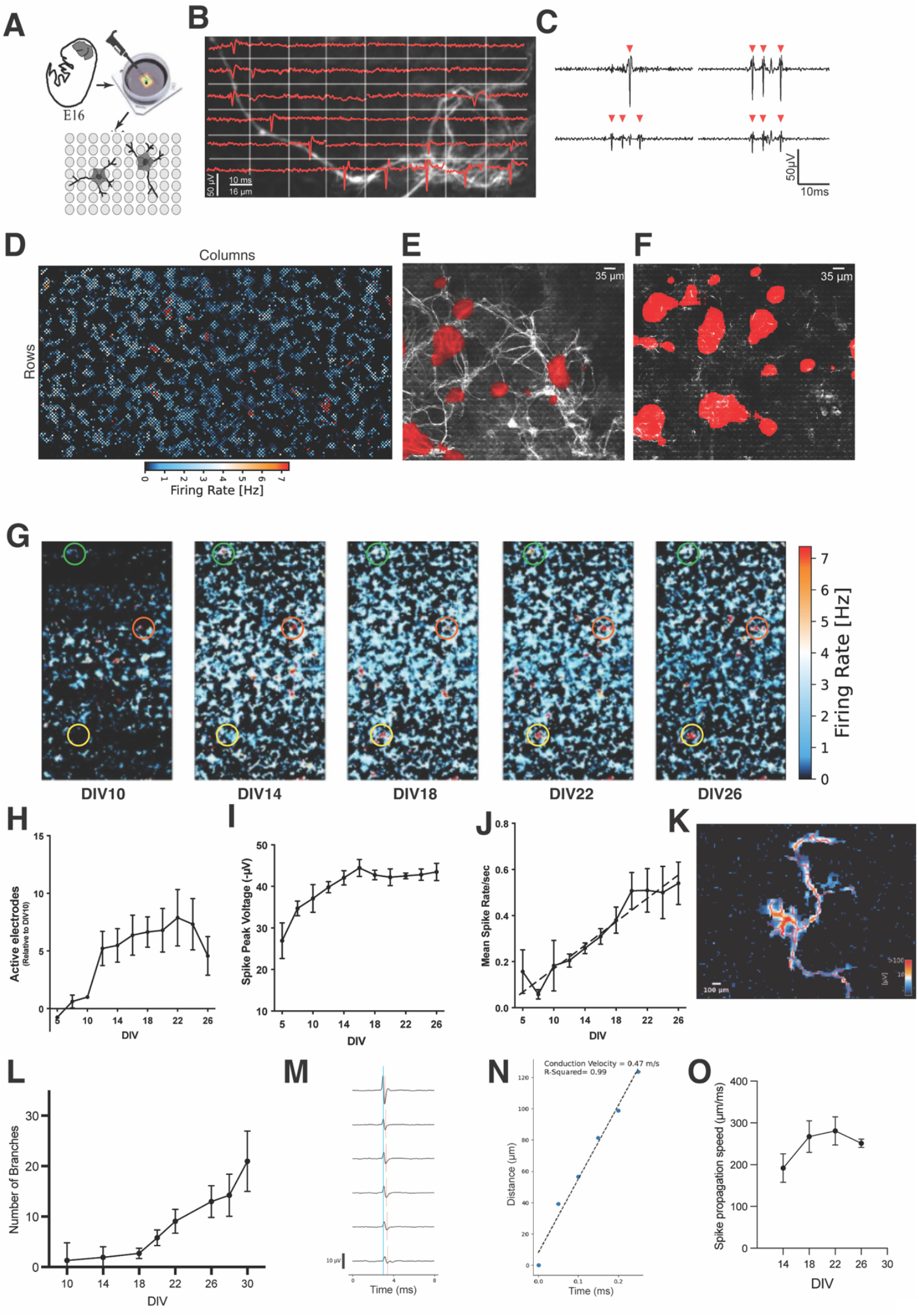
Characterization of normal electrical activity development. (A) Schematic diagram of the preparation of E16.5 mouse cortical neurons for culture on the MEA. (B) Overlay of immunostaining of MAP2 (white), traces from spontaneous recordings (red lines) and the boundaries of the 48 individual electrodes measured (grey squares). Wildtype culture, DIV26. (C) Examples of events scored as spikes (red triangles) recorded from four different electrodes. (D) A heatmap of the firing rate of a wildtype DIV 18 culture during a 1-minute period. (E) Overlay of designated hotspots (red) with immunostaining of MAP2 (white) performed after the recording session. (F) Overlay of hotspots and immunostaining of Ankyrin G (white). (G) Successive heatmaps of the firing rate of a single MEA culture made over a two-week period (DIV as indicated). Green circles illustrate a single group of electrodes that changed from active to inactive during the culture period. Red circles show a spot that became active and remained so at all subsequent DIVs. Yellow circles indicate a spot that became active only at late DIVs. (H) The number of active electrodes in the wildtype culture normalized to the number found at DIV10. (I) Mean spike amplitude (−µV) of wildtype cultures as a function of DIV. (J) The mean spike rate of wildtype cultures as a function of DIV. The dashed line is the least squares fit of the points. (K) An example of the tracking of axons and their branches based on the timing of their spiking activity (see text for details). (L) The number of axon branches of neurons in a wildtype culture expressed as a function of time in culture. (M) Spikes recorded from electrodes adjacent to the reference electrode. Note the slight delay in the peak depolarization from one electrode to its neighbor. (N) Spike peak time plotted as a function of distance from the reference electrode. The slope of the line represents the least squares fit of the speed of propagation. (O) The speed of spike propagation in a wildtype culture as a function of DIV. For Panels H-J. L, and O, n=10 independent WT cultures; error bars represent the standard error of the mean.

Monitoring the activity of the culture, we defined a voltage shift as a “spike” when the peak potential shift exceeded six times the standard deviation of the average baseline signal (Fig. 1C, red triangles). We sequentially sampled all 26,400 electrodes of the MEA during a 30-minute recording session and obtained the spike rate of each electrode, which we displayed as a heat map of the array (e.g., Fig. 1D). For each session, we identified the 1,024 most active electrodes to use for further analysis. We found these to be scattered across the full extent of the array (Fig. 1D), suggesting that clumping of cells or processes was minimal in our preparations. Superimposition of the location of the most active electrodes on a corresponding image of the immunostained culture revealed that they were located predominantly at the cell body (MAP2 immunostain – Fig. 1E), with a bias toward the location of the axon initial segment (ankyrin G immunostain – Fig. 1F).

Recording sessions were conducted over a period of 26 days, allowing us to track the total number of active electrodes as a function of time in culture. The images in Fig. 1G are heat maps of sequential recordings from a single array over a 2 ½ week period from DIV10 through DIV26. Earlier time points are not shown because there was little electrical activity evident before DIV10. Between DIV10 and DIV14, however, the number of active electrodes increased significantly, after which there was little additional change (Fig. 1G, H), a pattern seen by others (e.g., ^28,29,30^). Yet, while the number of active electrodes did not change, the relative activity at any one electrode could vary considerably. Some sites began with nearly no activity then gradually developed a high spiking rate (Fig. 1G, yellow circles). Others oscillated between no activity in certain time windows to high activity in others only to become quiet again at later times in culture (Fig. 1G, green and orange circles). Other electrophysiological properties of the culture appeared to develop on independent time schedules. For example, unlike the near saltatory increase in the number of active electrodes between DIV10 and DIV14, the average peak spike voltage of a given element increased steadily until DIV16 after which it plateaued for the rest of the culture period (Fig. 1I). The mean spike rate showed yet a third pattern, increasing nearly linearly throughout the entire 3–4-week culture period (Fig 1J). Heat maps illustrating these properties provide a sense of how these different properties mature (Supp. Fig. 1).

There was also a significant increase in the number of active branches of each axon. To follow individual action potentials as they left their site of initiation and traveled across the culture substrate, we identified the 15 most active electrodes and recorded from a 9×9 matrix of 81 electrodes surrounding the 15. We then traced spike events as they moved across the MEA (Fig. 1K, Supp movie 1)^31^. These data also allowed us to count the number of active axonal branches. During the first two weeks in culture most axons displayed an average of 3-4 branches, but after DIV18 the number of branches increased nearly four-fold (Fig. 1L). The recording data were also used to estimate axonal conduction speed (Fig. 1M, N), which trended upward until DIV16 after which it appeared to stabilize (black symbols, Fig. 1O).

### Network activity of the array begins with large scales bursts and matures to more local activity

Viewed horizontally, the MEA consists of 110 rows of electrodes arrayed in 220 columns. To display network activity, we numbered the 1,024 most active electrodes starting at the top of the first column (top left in Fig. 2A) then proceeded through each of the 220 columns to the bottom right of the array. In this example, we have used a color gradient to illustrate the order in which the electrodes were sampled. These electrodes were then displayed in order along one axis, with time in seconds plotted on the other axis resulting in a graph such as the one shown in Figure 2B. Note how some electrodes display a near constant level of spiking. These elements appear as a vertical band in the raster plot (Fig. 2B, magenta arrows); others were far less active (Fig. 2B, green arrows). The network also showed evidence of coordinated bursting activity (horizontal bands, Fig. 2B, orange arrows). Some bursts were quite strong and involved virtually every electrode (Fig. 2B, gray arrows).

**Figure 2.**
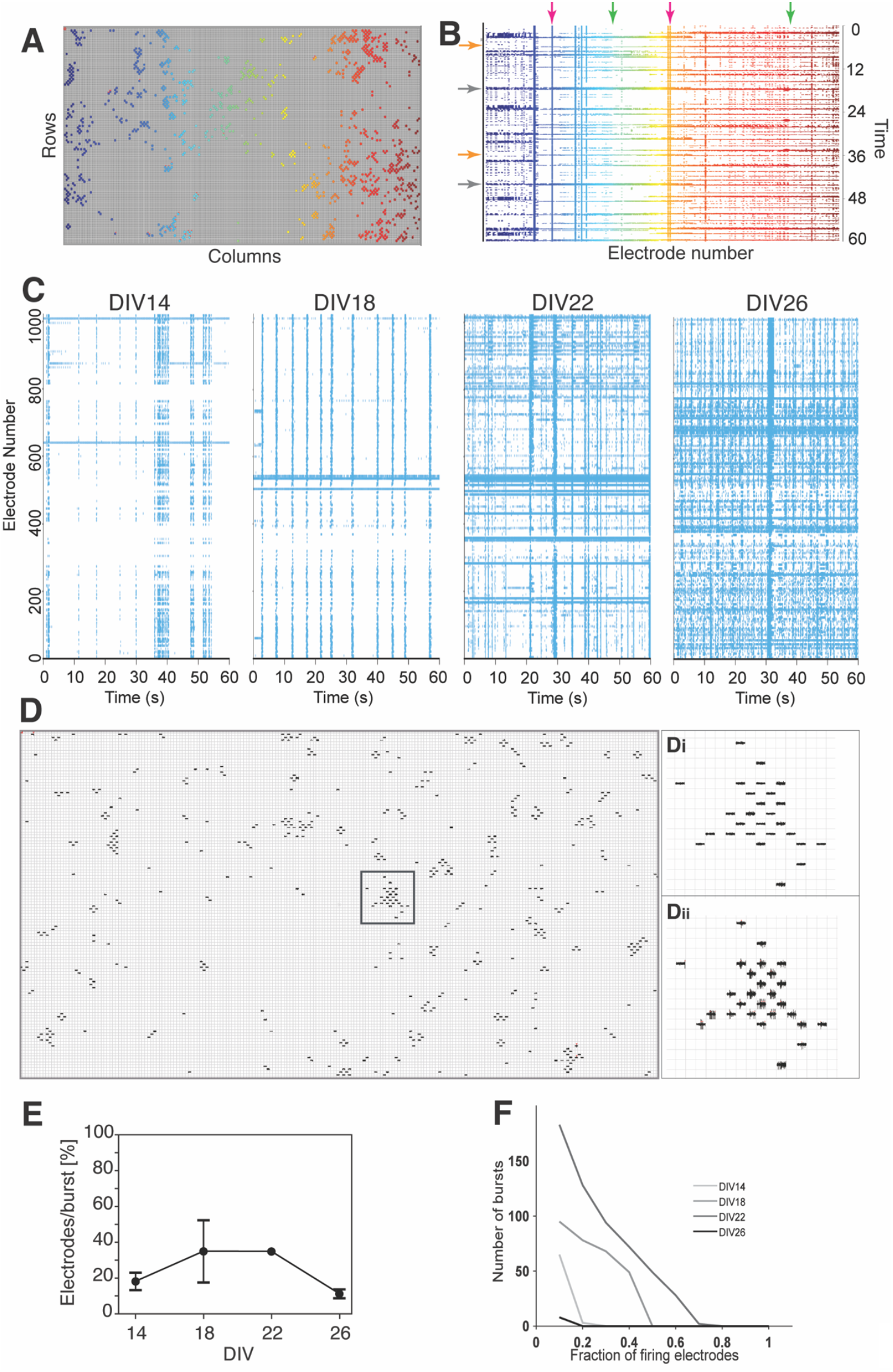
Network activity development. (A) The position of all active electrodes on a typical MEA based on a 5-min recording of a wildtype culture done at DIV18. The active electrodes were numbered, starting at the upper left and proceeding, column by column to the lower right. A color scale was used to reflect the number assigned to each electrode. (B) Raster plot showing the activity of each of the electrodes (colored as in (A)) during a representative 60 second time window of the 5-minute recording session. Each dot represents a detected spike at the electrode. Red and green arrows indicate constitutively active electrode sites. Gray arrows represent bursts (simultaneous activity) involving a high percentage of sites. Orange arrows represent bursts involving a lower percentage. (C) Raster plots of a single wild type culture recorded at 4-day intervals beginning at DIV14. (note: plots are rotated 90 degrees, relative to the plot in panel B). (D) Example of bursting activity on the array. In the diagram to the left each grid square contains the activity of that electrode during a 5 ms recording interval. Di and Dii are enlargements of the grid squares indicated by the black box taken at two different times during the recording session. Di illustrates a 5 ms quiescent period; Dii represents the same electrodes during a burst. (E) The fraction of total number of active electrodes participating in a single burst at different DIV. (error bars = SEM) (F) The number of bursts as a function of the fraction of the total number of electrodes involved in that burst. n=10 independent cultures for panels E and F.

Using raster plots such as this we followed the development of the activity of the cultures over time. At first (DIV14), coordinated bursting activity predominated (Fig. 2C – note that in this representation the axes have been reversed from panel B; bursts are now vertical; active sites are horizontal). Similar patterns have also been observed by others (e.g., ^29,30,32^). Bursting activity matured with time in culture, such that more and more units were involved in each burst (Fig. 1D-F). After DIV22, however, the number of electrodes participating in each bursting event declined sharply (Fig. 2E), as did the fraction of the total electrodes that were involved in any one burst (Fig. 2F). This likely reflects the establishment of local circuits that fire independently of the other parts of the network.

### Altering the levels of APP in mature cultures affects individual units more than the network

Aging and Alzheimer’s disease are closely correlated with increased levels of APP in the brain. We have previously shown that APP increases with activity and such increases alter the structure of the axon initial segment (AIS), which is predicted to quiet a neuron^33,34^. This prediction has been validated with calcium imaging that the neurons carrying APP_Swe_^35,36^. These findings led us to probe the functional consequences of acute changes in APP levels using the high resolution afforded by the MEAs. We infected DIV10 cultures with a lentivirus (lentiAPP) that expressed full length APP_Swe_ plus an mCherry marker that allowed us to identify the infected cells (Fig. 3A). On DIV18 we identified the most active neurons then applied bi-phasic 200 µsec pulses of increasing voltage spaced one second apart. Recordings from the surrounding electrodes were then used to determine whether the stimulus had elicited a response. For each neuron and condition we performed 20 replicates. (see Methods for additional details). Increasing the applied voltage to uninfected cells increased the probability of firing, with a half maximal response seen at ∼35 mV (Fig. 3B, black symbols). By contrast, the lentiAPP cultures required nearly 80 mV to achieve the same half-maximal response (Fig. 3B, purple symbols). Consistent with the reduced probability of firing, the average rate of spontaneous spiking was reduced in lentiAPP cultures. This effect became more pronounced as the culture matured (Fig. 3C). Other properties of the individual units were unaffected. The amplitude of the average spike event was largely unchanged (Fig. 3D) and the number of active electrodes was not reduced (Fig. 3E).

**Figure 3.**
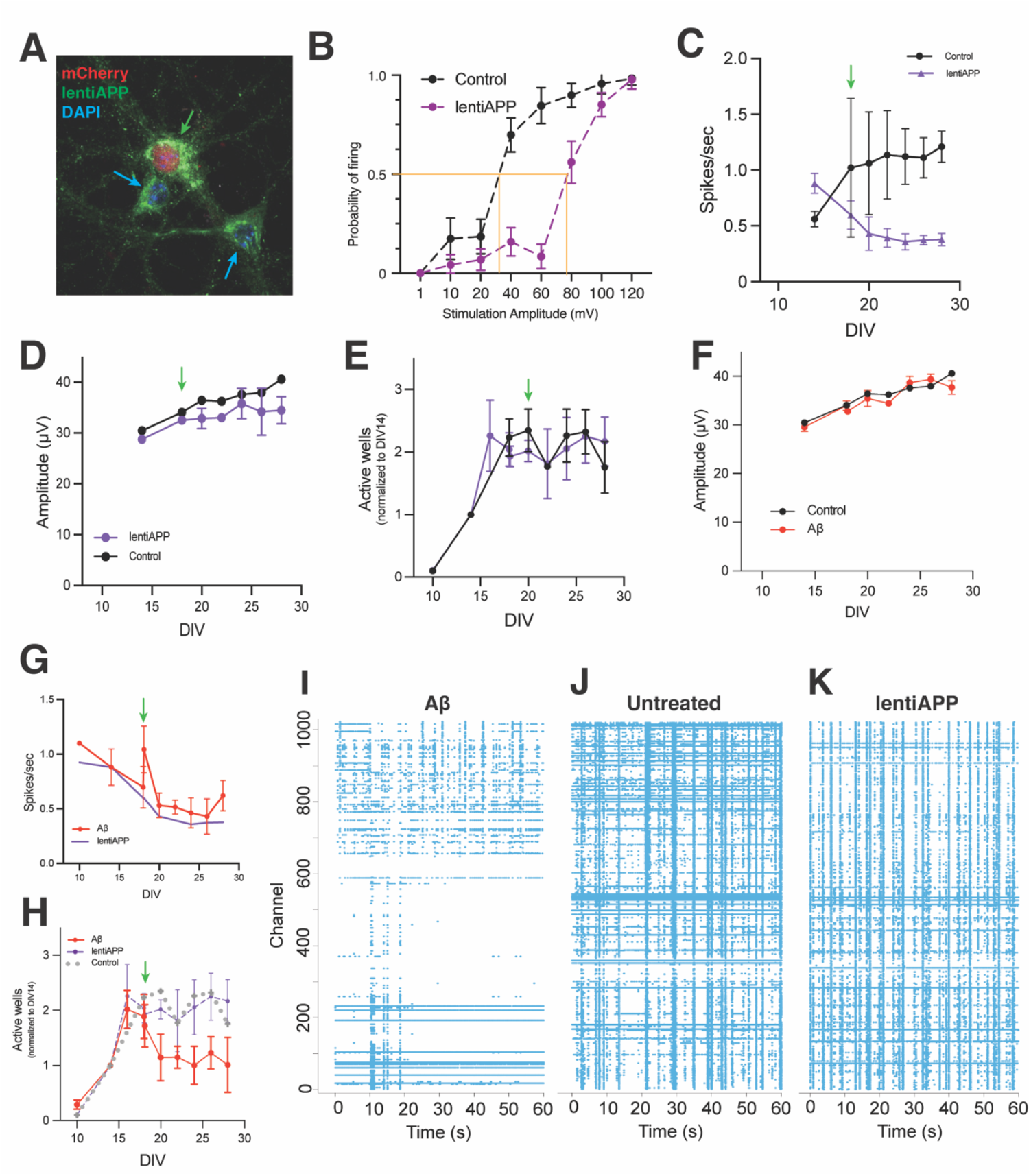
Effects of lentivirus expressing APP_Swe_ and Aβ treatment at DIV18. (A) Three neurons from a lentiAPP infected culture. Nuclear mCherry (red) identifies an infected cell. Immunostaining for APP (green) confirms that APP levels increase in infected (green arrow) compared to uninfected (blue arrow) cells. (B) Probability of a site firing after stimulation with different voltages in wildtype (black) and lentiAPP (purple) cultures. Orange lines indicate the voltage at which 50% firing probability is reached. (C) Firing rate of lentiAPP and control cultures as a function of DIV. (D) Spike amplitude as a function of DIV for lentiAPP and controls. (E) The number of active electrodes in control and lentiAPP cultures (normalized to DIV14). (F) Amplitude of spikes in control and Aβ oligomer treated culture. (G) The effect of Aβ oligomers on spike rate. The purple line is a reproduction of the lentiAPP data from (C) included for reference. (H) The effect of Aβ oligomer treatment on the number of active electrodes. The dashed gray line (wild type) and purple graph (lentiAPP) are a reproduction of the data from panel E included for reference. For panels C-H, the green arrows indicate the DIV of lentiAPP infection or Aβ oligomer addition. In all graphs error bars represent the standard error of the mean. (I-K) Representative raster plots recorded on DIV22 after treatment with Aβ oligomer, control, or lentiAPP on DIV18. n=10 independent wildtype cultures and n=8 independent lentiAPP cultures shown in panels B-G.

Although we had previously shown that exogenously applied Aβ (fibrillar or oligomeric) had no effect on the structure or location of the AIS, we felt it was important to test whether Aβ was mediating any of the observed lentiAPP effects. We applied 1 µM Aβ oligomers to mature cultures (DIV18) and measured their responses. The oligomers had no effect on spike amplitude (Fig. 3F) but the spike rate, after a transient increase in the hour following Aβ addition (p < 0.01), decreased and closely tracked the reduced rate seen in the lentiAPP cultures (Fig. 3G). On its own, this finding would be consistent with Aβ mediating the APP effects, but other metrics offered a different perspective. The number of active electrodes was unchanged by lentiAPP, yet adding Aβ oligomers reduced the number active electrodes by half (Fig. 3H, p < 0.001). Infection with lentiAPP had little obvious effect on the network activity seen in the raster plots; addition of Aβ, by contrast, quieted entire regions of the array leaving other regions largely unaffected (Fig. 3I-K). Interestingly, the development of axonal branching was similar to that found in WT cultures (Supp Fig. 2).

### In less mature cultures APP retards or blocks the development of the network

The development of activity in the cultures progressed through a period of synchronous activity (bursting) then matured to a pattern characterized by more local interactions (Fig. 2I). We noted that four days after DIV18 lentiAPP infection, cultures appeared to regress towards a DIV14 pattern of increased bursting. This led us to ask whether infecting cultures earlier in their maturation process might lead to a similar regression. When we infected cultures with lentiAPP on DIV10 instead of DIV18, we found that the development of the culture was profoundly blocked. The number of active electrodes increased slightly, but the dramatic increase found in the untreated cultures was blunted (p < 0.001) (Fig. 4A). A similar pattern was seen in the development of spike amplitude. The effect was less dramatic, but shortly after lentiAPP infection, the increase in amplitude found in the wild type was blocked (Fig. 4B). For both measures, the block was maintained for 2 ½ to 3 weeks following infection. The developmental block, however, was not complete. The spike rate followed a bi-phasic response during which it first dropped (p < 0.001 for DIV 16 and DIV 18 compared to control), then between DIV18 and DIV20 rapidly increased before finally plateauing slightly below the rate found in untreated cultures (Fig. 4C).

**Figure 4.**
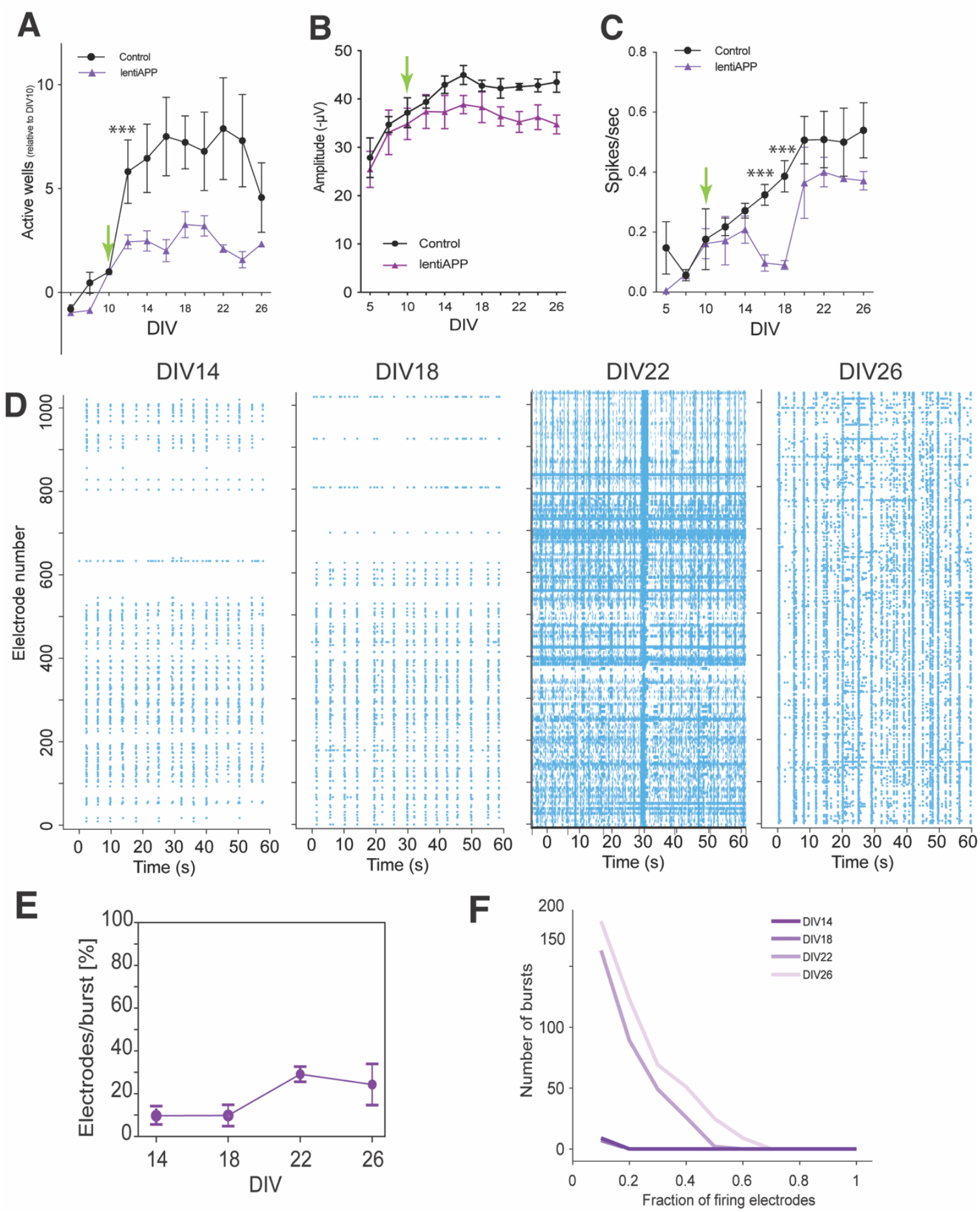
APP_Swe_ lentivirus treatment on DIV10. (A) The number of active electrodes after infection with lentiAPP normalized to DIV10. In this and subsequent plots, the green arrows indicate the DIV when lentivirus was added to the culture, and the purple line represents the control data from Figure 1 replotted for reference. (error bars = SEM, *** p < 0.001 relative to control). (B) The average spike amplitude as a function of DIV after treatment with lentiAPP on DIV10. Error bars = SEM. (C) The mean firing rate recorded after infection with lentiAPP on DIV10 plotted as a function of DIV. (error bars = SEM, *** p < 0.001 relative to control) (D) Representative raster plots of a single lentiAPP culture recorded every four days beginning at DIV14. (E) The percentage of active sites involved in a burst as a function of DIV. n=8 independent cultures. Error bars represent the standard error of the mean) (F) Number of bursts plotted as a function of the fraction of the total number of electrodes involved in that burst. n=8 independent cultures for panels A, C, E, F.

The block in the development was also seen in the raster plots. As early as DIV14, the pattern of activity on the array was reduced relative to untreated wild type cultures (Fig. 4D, compare with Fig. 2C) and this reduction continued through DIV18. The pattern of firing with time in culture was also notably different from that seen in untreated cultures. On DIV14, four days after infection, bursting activity was present, but its frequency trailed that seen in age-matched untreated cultures. By DIV18 and beyond the immature pattern persisted, and did not mature with additional time in culture (Fig. 4D). Note in particular the absence of the inter-burst activity that characterized the more mature untreated control cultures (Fig. 2C). Plotting the number of bursts as a function of the percentage of active electrodes in a burst offers a more quantitative picture of this lack of developmental change (Fig. 4E-F). The appearance of coordinated bursting was slow to develop in the lentiAPP cultures such that the pattern at DIV26 more closely resembled younger untreated cultures (Fig. 2F). Consistent with the biphasic effects of lentiAPP on spike rate (Fig. 4C), there was a transient increase in activity visible in the DIV22 raster plot. Despite this, by DIV26, the lentiAPP cultures returned to their immature state with a noticeable absence of continuously active recording sites (horizontal bands in the DIV26 raster plot in Fig. 4D). We also observed fewer axonal branches on the active neurons (Supp Fig. 2) compared to untreated and Aβ-treated cultures.

To ask whether the Aβ peptide, rather than full-length APP, was driving these phenomena, we exposed sister cultures to Aβ oligomers at DIV14, adding fresh Aβ every four days to maintain its presence. The Aβ-treated MEAs differed from both untreated and lentiAPP cultures. While lentiAPP infection at DIV10 virtually halted the increase in the number of active sites, Aβ exposure led to only a modest decrease from controls (Fig. 5A). The spike amplitude (Fig. 5B) drifted lower but was largely identical to untreated cultures. The most unexpected result came from our analysis of the average spike rate. While lentiAPP infection at DIV10 reduced spike frequency, Aβ addition markedly increased it (Fig. 5C). Curiously, Aβ also appeared to block the normal increase in conduction speed observed in untreated cultures (Supp. Fig. 3). Aβ also had a strong effect on network activity. Similar to earlier findings^37-40^, one hour after Aβ addition, we observed an increase in firing frequency (Fig. 3G). Compared to the effects of lentiAPP, Aβ induced almost no bursting activity (Fig. 5D) and there was enormous variation in the activity pattern of the Aβ-treated cultures. We observed only a small number of bursts with less than 30% of the active electrodes participating in the days immediately following Aβ addition (Fig. 5E), but even this modest level of coordinated activity disappeared by DIV22 (Fig. 5E-F). In addition, the number of active sites participating in any one burst was reduced from controls (p < 0.05). By contrast, sites with continuous activity were more common than in lentiAPP cultures, though not as frequent as in untreated cultures. We performed 6E10 immunostaining of sister DIV18 coverslip cultures to learn if there were structural correlates to this odd and variable behavior. The staining pattern revealed small domains of neuropil where synapsin-I staining was virtually absent. In these regions, the MAP2^+^ neurites and spine-like protuberances were coated with small puncta of 6E10 immunostaining (Fig. 5G). The staining suggests that there had been a loss of morphological synapses, which would fit with the loss of network activity, and suggests that Aβ oligomers act in part by breaking both structural and functional synaptic connections.

**Figure 5.**
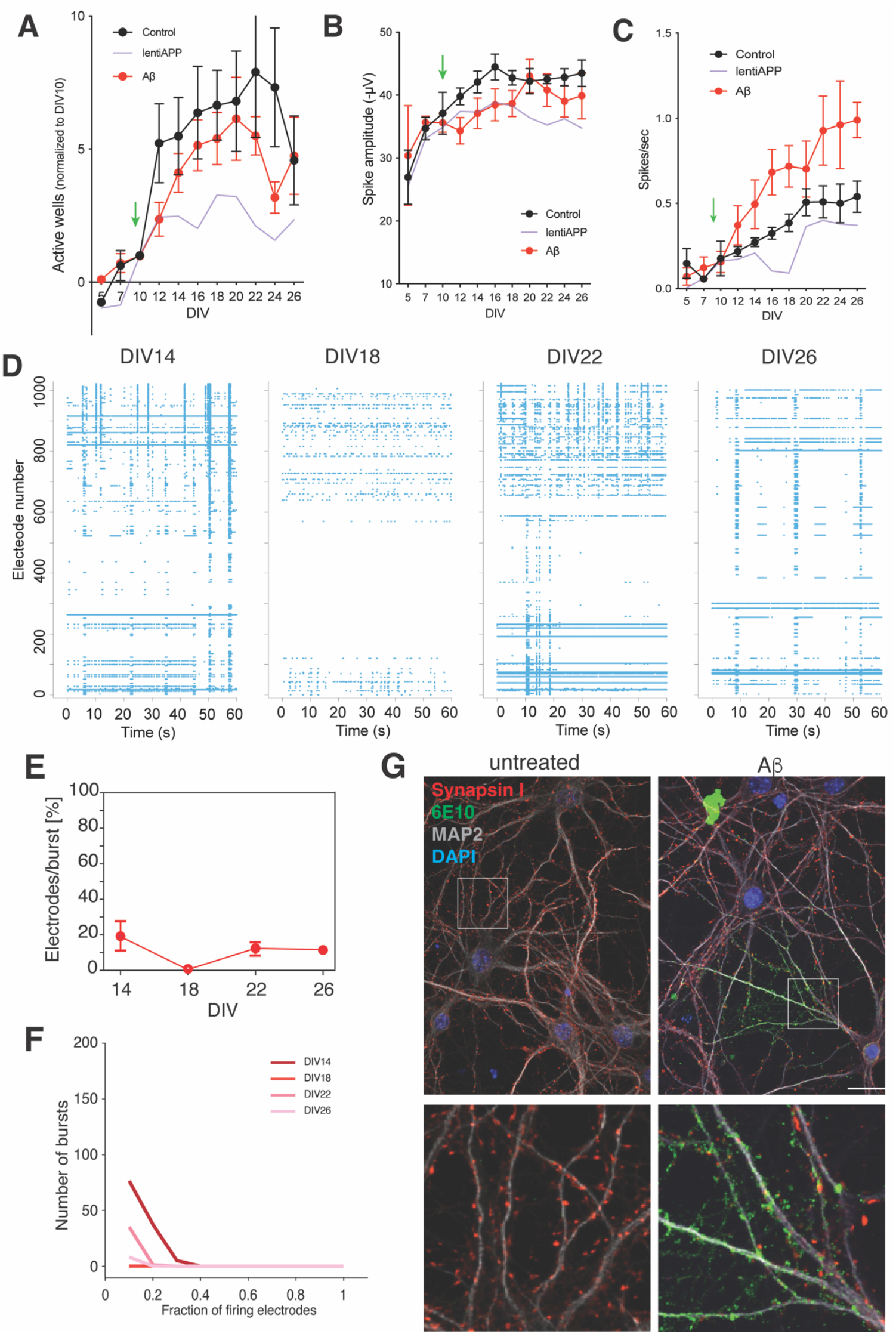
Aβ oligomer addition at DIV10. (A) The number of active electrodes after treatment with Aβ oligomer normalized to the values at DIV10. In this and subsequent panels, the green arrows indicate the DIV when Aβ oligomers were first added to the culture. (B) The average spike amplitude as a function of DIV after treatment with Aβ oligomers at DIV10. (C) The firing rate of the active electrodes in a culture following treatment with Aβ oligomer at DIV10. In (A) – (C) the purple line represents the lentiAPP data from Figure 3 and the black points are data from Figure 1 replotted for reference. (D) Representative raster plots of a single culture treated with Aβ oligomer on DIV10 recorded every four days beginning at DIV14. (E) The percentage of active sites involved in a burst as a function of DIV. (F) Number of bursts plotted as a function of the fraction of the total number of electrodes involved in that burst. (G) Immunolabeling of a wildtype neuronal culture either untreated (left) or treated with 1 µM Aβ oligomer for 24 hours (right). Synapsin I (red), 6E10 (green), MAP2 (gray), and DAPI (blue). Scale bar = 20 µm. Areas in the white boxes are enlarged to illustrate how, in Aβ-treated cultures, there are areas where synapsin-I staining is missing and seemingly replaced by puncta of 6E10. n=5 independent Aβ cultures for panels A – F. In panels A-C and E, error bars represent the standard error of the mean.

The contrast between applying APP or Aβ to mature or developing cultures led us to ask what the consequences would be if the levels of APP were elevated throughout the culturing process. To achieve this, we cultured neurons from R1.40 transgenic mice, a model of human AD that carries a yeast artificial chromosome transgene with the entire human APP_Swe_ gene including introns and all cis regulatory elements within 30 kb of the coding sequence. In R1.40 cultures, the number of active electrodes was dramatically reduced (p<0.001) compared to untreated cultures but was similar to the number found in DIV10-infected lentiAPP cultures (Fig. 6A). By contrast, spike amplitude developed in nearly the same way as it did in untreated cultures as opposed to the reduced values seen in the lentiAPP cultures (Fig. 6B). Indeed, at the earliest time points (DIV5 and DIV7) the R1.40 spike amplitude was significantly greater than untreated cultures. This precocious developmental pattern was also found in the emergence of the spike rate (Fig. 6C).

**Figure 6.**
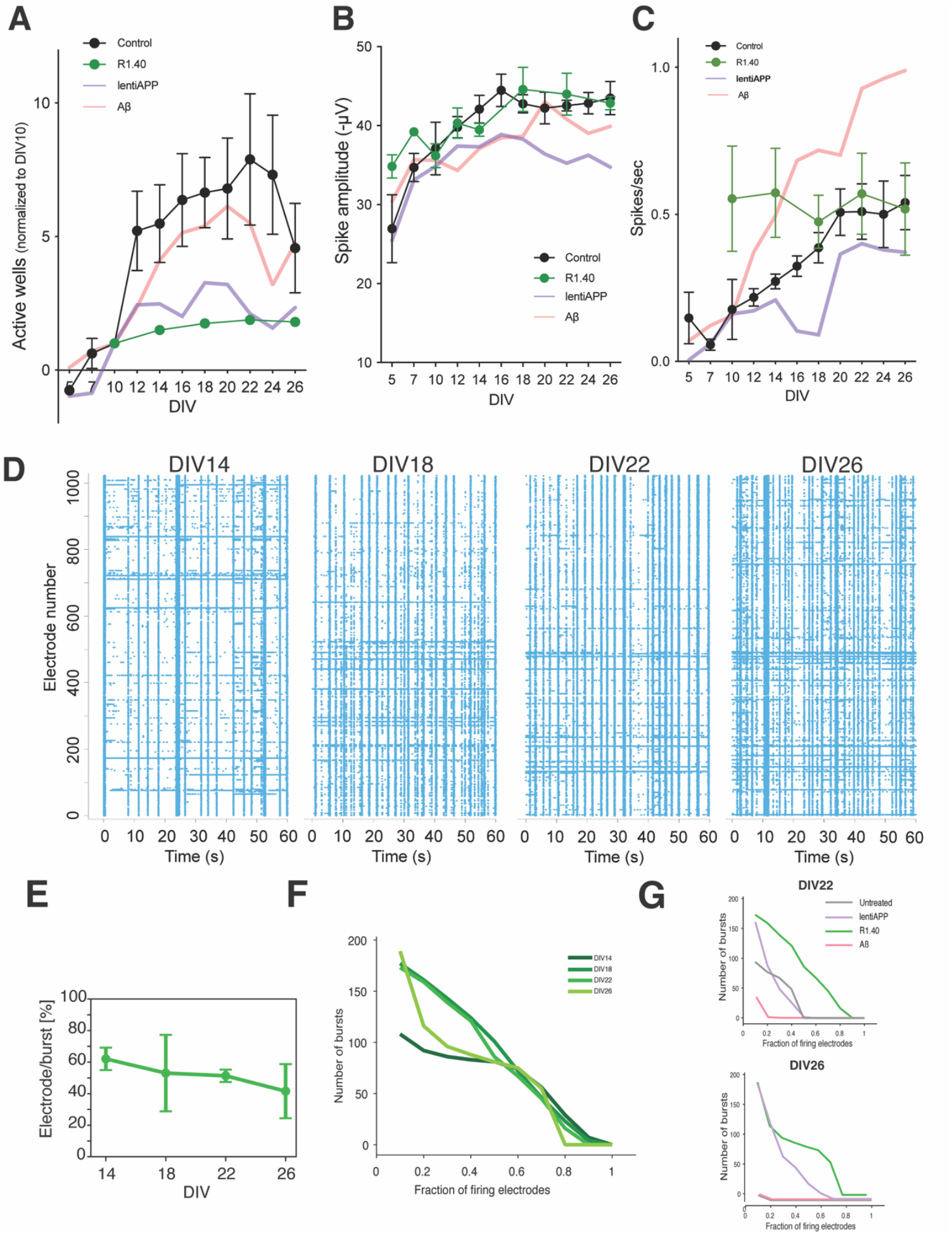
Behavior of R1.40 neurons in MEA cultures. (A) The number of active electrodes in R1.40 cultures normalized to the values at DIV10. (B) The average spike amplitude in R1.40 cultures as a function of DIV. (error bars = SEM) (C) The firing rate of the active electrodes in R1.40 cultures (error bars = SEM). In panels (A) – (C), the purple line is a re-plotting of the lentiAPP data from Figure 3; the pink line is a re-plotting of the Aβ oligomer data from Figure 5 and the black points are the control data from Figure 1 included for reference. (D) Representative raster plots of R1.40 cultures at different DIVs. (E) The average percentage of active sites involved in a burst as a function of DIV. (F) Number of bursts plotted as a function of the fraction of the total number of electrodes involved in that burst. (G) Re-plotting of the data from Figs. 2F, 4F, 5F, 6F to allow the comparison of the four types of cultures with respect to the number of bursts as a function of the fraction of total electrodes active at DIV 22 and DIV 26. n=6 independent R1.40 cultures for panels A – F. In panels A-C and E, error bars represent the standard error of the mean.

The network activity of the cultures was, in general, distinct from untreated and Aβ treatments but behaved slightly similar to lentiAPP infected cultures. At DIV14 R1.40 cultures had already established a pattern of burst and inter-burst activity that resembled more mature untreated cultures, yet there appeared to be little further development beyond this (Fig. 6D). This can be visualized by viewing the change in the fraction of sites contributing to a burst (Fig. 6E-F). Whereas wild type cultures have a transient period of coordinated bursting activity, they soon lose this feature. R1.40 cultures develop earlier but seem locked in their pattern of bursting (Fig. 6F). This was also corroborated by axonal branching data where we found fewer branches on the active sites, similar to the lentiAPP cultures (Supp Fig. 2). This developmental block is reminiscent of, but more extensive than, that seen with DIV10 lentiAPP. The plots in Figure 6F are reproductions of the data from earlier figures to allow a head-to-head comparison of the network activity of the four types of cultures as they mature. They highlight the extent to which the R1.40 cultures are retarded in their maturation and suggest that DIV10 lentiAPP infections come close to but do not entirely replicate this block.

## Discussion

We have exploited the possibilities afforded by CMOS technology to monitor cortical neuronal cultures with 26,400 closely spaced contiguous electrode tiles. The resulting electrical view of cultures spans considerable distances (8 × 10^6^ µm^2^) at very high resolution. Despite the complexity of the electronics and their surfaces, there was no tendency for cells to clump or for neurites to track along the interfaces among the surface tiles. Further, as we recorded directly in the CO_2_ incubator, we could easily perform longitudinal studies, tracking a single culture over several weeks, with minimal stress on the cells.

We found that the electrical features of the cultures developed at different rates. Neurons were largely quiet during early days in culture, then became electrically active between DIV10 and DIV14. This timing is consistent with the reports from other laboratories^29,30,41-44^. Following this, there was little further increase. Two features of this MEA data bear noting. The first is that most of the active electrodes that appear in our cultures do not reflect neuronal cell bodies, but rather reflect the activity of their axons. The second is that not all processes that are present in the culture are active. Both features can be seen in Figure 1 (Fig. 1B, E, F). Neurons are post-mitotic in vivo and in vitro so their numbers do not increase after they are plated. By contrast, structural features of the cultures – dendritic and axonal branches – expand continuously. As the number of active electrodes remains constant after DIV14 (Fig. 1A), the suggestion is that the constant number of neurons puts a ceiling on the level of activity that can be achieved. Our finding that activity recorded at individual electrodes waxes and wanes with time in culture is consistent with this idea.

Viewed as a whole, the activity of the cultures reflects two defining features – constitutive activity of individual sites and coordinate activity of the entire ensemble (bursting). Bursting develops first, as reported by others^29,30,45,46^. When viewing the behavior of the entire culture we find that over the first three weeks, the number of electrodes that are active during a burst increases before dropping precipitously after DIV22 (Fig. 2E, F). The impression is that after three weeks local circuits begin to take the place of pan-culture activities as has been proposed by others^47,48^.

Against this detailed picture of the behavior of the neuronal ensembles, the responses of the cultures to increasing the levels of the amyloid precursor protein (APP) are highly informative. APP, true to its name, has been largely studied as a source of the β-amyloid peptide. Yet our results encourage a broader view of the functions of full length APP itself. Given our earlier findings^27^, it was gratifying to find that lentiAPP increased the voltage needed to fire a neuron (Fig. 3B) and thus decreased the average firing rate observed across the entire network (Fig. 3C). These results are consistent with those of Meza et al^26^ who observed that the firing rate of neurons was negatively related to the distance between the cell soma and the proximal end of AIS. The broader network of cells, however, proved resilient to altering the levels of APP in these relatively mature cultures as the number of active electrodes was largely unaffected (Fig. 3E). Axonal branching data combined with the early maturation of the networks suggest that APP overexpression primarily works at a single neuron level by reducing spike probability and blocking additional connections with other neurons. This leads to fewer but more inclusive network-wide bursts. We propose that this precludes the cultures from forming smaller local networks such as those seen in untreated cultures.

By contrast, infecting the culture with lentiAPP at DIV10 had a dramatic impact both on the behavior of the individual units (spike amplitude and spikes per second) as well as the activity of the larger network of cells (number of active electrodes and the blocked maturation of mature activity patterns). Coupled with the relative lack of impact of exactly the same manipulations performed at DIV18, the data suggest that two different processes are at work. The first is the homeostatic effect of APP on a mature neuron. Through changes in the AIS, APP quiets an active neuron and thereby functions as a feedback governor to protect it from over-heating. The DIV10 data, extended by the findings in the R1.40 cultures, suggest that in addition to this homeostatic role, APP has a prominent role to play in neuronal maturation. As the AIS is found in cultures as early as DIV4, it is conceivable that the network develops more slowly (or not at all) when the activity of its individual units is blunted, but other explanations are possible.

The Aβ data adds additional complexity to the situation. The oligomeric aggregates of Aβ affect the cultures in ways that are distinctly different from those found in lentiAPP or R1.40 cultures. Addition of Aβ to mature (DIV18) cultures reduced the spike rate in much the same way as lentiAPP but reduced the number of active electrodes by half compared to lentiAPP and control cultures (Fig. 2H). The differences with lentiAPP were more apparent when Aβ treatment was begun at DIV10. Unlike the profound reduction in the number active electrodes found after lentiAPP, Aβ addition had only a minimal effect. Further, while lentiAPP reduced the spike rate, Aβ significantly increased it. Similarly, the response of the network to Aβ was completely unlike its response to lentiAPP. Aβ almost completely suppressed bursting activity while lentiAPP appeared to block the ability of the network to stop bursting. Notably, adding oligomers to mature cultures does not alter the structure of the AIS^35^. Taken together, the data suggest that APP and Aβ produce their effects by different mechanisms. Our interpretation of the overall findings is that early on, APP retards nerve cell development, while in the adult it dampens neuronal excitability. By contrast, at all ages, Aβ appears to break existing synaptic connections, a prediction consistent with the patches of 6E10 immunostaining that are devoid of synapsin I immunostaining (Fig. 5I). Similar colocalization pattern of Aβ peptides and synapses has also been demonstrated recently^49,50^.

The data supports a model where APP acts on individual developing neurons, making them less likely to spike but leaving existing connections intact. By contrast, Aβ works at the network level. By making the individual neurons hyper-active but reducing the synaptic connections between them it leads to an overall lower firing level within the network. The branching data suggests that neurons in Aβ cultures can still create many functional branches whereas over-expressing APP early in development stunts this axonal growth. This is consistent with our finding that later addition of APP does not have the same effect on axonal branching as earlier addition, and therefore does not alter network connections once they have formed.

The findings have important implications for our understanding of Alzheimer’s disease. First, the lack of concordance between the actions of APP and exogenous Aβ leads to the suggestion that the clinical symptoms of the disease may represent a composite response of the AD brain to both biological agents. Many studies have concluded that the location of amyloid plaques and the cellular structural changes related to Alzheimer’s disease^51,52^ are a poor predictor of the cognitive deficiencies experienced by any one person. In light of our current findings, we note that analyses of the location and concentrations of full-length APP are rarely reported, and submit that this information would be a valuable part of a more complete picture of AD pathogenesis.

Second, since the editorial published by Katzman in 1976 ^53^, it has been widely accepted that early onset familial forms of AD, caused by *APP, PSEN1*, and *PSEN2* mutations represent the same disease process as the more common sporadic forms. Our results suggest that this assumption may not be warranted. Late infection with lentiAPP would be a model of sporadic AD, while R1.40 neurons would represent familial/genetic forms. R1.40 neurons display cell-autonomous alterations in their neurophysiology that contrast starkly with lentiAPP or control cultures. Higher spike amplitudes and spike rates appear precociously in R1.40 cultures as does the bursting pattern. The cultures then appear blocked in their development and never lose the more immature coordinated bursting that disappears in controls. This suggests that genetic forms of AD may prime a person towards dementia onset through a different set of biological mechanisms than the late-life elevations of APP and Aβ seen in the far more common sporadic form of Alzheimer’s.

## Materials and Methods

### Primary culture

The culture surface of the MEA chip was coated with poly-D-lysine (PDL, 0.5 mg/ml) in 37°C overnight. The PDL was rinsed off with water followed by an additional coating with 50µg/µL Matrigel (Sigma L2020) for 30 minutes. Wild type E16.5 C57BL/6J embryos were harvested from gravid females. After cervical dislocation, their cerebral cortices were isolated and stored at 4°C in PBS supplemented with 1 mM glucose. Cortices were separated into small pieces using forceps and treated with 0.25% trypsin-EDTA (Sigma) for 10 min at 37°C. The tissue was then washed in 10% FBS in DMEM to inactivate the trypsin, then washed in Neurobasal medium (Invitrogen), and triturated to produce a single-cell suspension. For cultures on glass, poly-L-lysine-coated coverslips were placed in the bottom of 24-well plates and cells added at a density of 48,000 cells/well. For MEA cultures, we used 50 µl of a 1.6×10^6^ cells/ml suspension to plate 80,000 cells total in the culture chamber (see below). The medium to maintain the neurons in both conditions was Neurobasal medium supplemented with 2% B27 (Invitrogen), 1% Glutamax (Invitrogen), and 10,000 U/ml penicillin/streptomycin (Invitrogen). Cultures were maintained at 37°C in a humidified atmosphere of 5% CO_2_. Every 5 d, the culture medium was refreshed by replacing half of the old medium. For APP overexpressing cultures, the cultures were infected with Lentivirus FUGW-APP_Swe_ (Addgene190804) produced by Dr. Ronald Hart^35^. For Aβ cultures, Aβ oligomers were prepared as described^54^. Briefly, the 1,1,1,3,3,3-Hexafluoro-2-Propanol (HFIP) pretreated Aβ^1-42^ (rPeptide, A-1163-1) were speed-vacuumed at 800 g at room temperature, then resuspend to obtain a final concentration of 100 µM. The resulting suspension was sonicated in a room water bath for 10 minutes then aliquoted into polypropylene vials, sealed and stored at –80 ºC. The freshly prepared oligomers were added to the cultures at a final concentration of 1 µM. Medium was refreshed every 4 days after DIV14 by removing half of the medium from each culture and adding back an equal volume of complete neurobasal medium. Freshly made Aβ oligomers were added with each feeding. The experimental flow is diagrammed in Fig. 1A.

### Electrophysiology

The electrical activity of the cultures was recorded using a single well CMOS based HD-MEA from Maxwell Biosystems Inc, ref. ^55^. The total array contains 26,400 tiles that function as both recording or stimulation sites. During fabrication the electrodes are coated with electrodeposited platinum black to decrease electrode impedance and improve the signal-to-noise characteristics. The CMOS technology allows the array to be constructed in such a way that the center-to-center distance of the tiles is only 17.5 µm. This enables a very high-resolution view of the electrical activity of the culture. While information can be retrieved from any electrode, the setup allows for simultaneous recording from only 1,024 of the total 26,400 electrodes at any one time at a sampling rate of 20 kHz.

#### Spike detection, sorting and analysis

Firing-rate based analysis was performed in Matlab (Mathworks, Inc.). The raw signals were filtered through a 300Hz high pass 2^nd^ order Butterworth filter. Spikes were detected using the level-crossing method where a change higher than 6 times the standard deviation of the recorded signal was considered a spike event. All analysis was performed on the firing rate based on these detected spikes. The recorded spikes were then sorted to detect individual units using the SpikingCircus ^56^ algorithm. All further analysis was performed using the sorted unit data. Spontaneous activity recording over the entire array. A gaussian convolution filter (sigma = 20 ms) was used to convert spike time into spike rate.

The MEA electrodes are arranged in a rectangular grid of 220 columns by 120 rows. We divided the array into two halves of 110 columns each. As the software allows a maximum of 1,024 electrodes to be recorded at any one time, we sampled the entire culture as follows. The first set of electrodes we recorded included the first 4 columns of each half of the array (columns 1-4 and 111-114). As each column contained 120 electrodes, each of the sets consisted of 960 electrodes which we monitored for 60 seconds. The second set of columns (5-8 and 115-118) was then recorded, and the process continued until all 27 full sets of electrodes had been recorded. The last two columns (119-120 and 219-220) were recorded as a final set. The data from all 28 sets were collected and analyzed as one in Matlab.

We calculated the total number of active electrodes (e.g., Fig. 1F) as the number of electrodes that spiked at least once during the 60-second recording. The mean spike rate of an electrode (e.g., Fig. 1G) was defined using only electrodes that showed at least one spike during the recording period. We summed the total number of spikes and divided this number by 60 to obtain the spike rate (spikes per second). We calculated the mean spike amplitude as the average amplitude across all recorded spikes from the active electrodes. We also observed a tendency for cultures to demonstrate bursting behavior. Bursting was defined as a local maxima of firing rate where most of the neurons in the network fired synchronously with an amplitude higher than 1.2 times the root mean square of the firing rate activity over the entire recording. The network activity assay was used to calculate the bursting frequency defined as number of bursts per second, inter-burst interval as the average duration between two network bursts and number of recorded electrodes per burst as the number of electrodes spiking at least once within 100 ms around the time of the burst. To track the bursting activity more accurately, we used the activity scan to identify the 1024 most active electrodes based on the firing rates. The spontaneous activity of the selected units was recorded for 5 minutes. For this analysis, bursting was defined as the synchronous activity of multiple neurons described in more detail in the Results. The spikes from all recorded neurons were plotted as a histogram with bin size of 10 ms, then smoothed using a gaussian kernel of width 300 ms.

#### Axonal propagation activity

The spontaneous activity of the 15 most active units of all 26,400 electrodes was recorded for 60 seconds. This was used to detect axons originating from the recorded 15 units. For each unit, spike triggered average was calculated for the rest of the electrodes to identify the axons^31^. The computation was performed using MaxLive software (Maxwell Biosystems, AG, Switzerland). Once the axons were identified the axonal length and conduction speed were calculated for each unit ^57^.

Axonal branching was defined as different branches of a propagating spike arising either from the soma or from the main axon. The branch was defined as a set of electrodes that form an axon and have a different spatial angle of propagation. The branching analysis and results were calculated using the MaxLive software (Maxwell Biosystems, AG, Switzerland).

#### Stimulation activity

The previously identified 15 units were stimulated using 20 bi-phasic pulses of 200 microseconds duration spaced 1 second apart. The amplitude of the pulses was from 2 mV to 240 mV peak to peak. Following stimulation, 1000 electrodes within 300 µm of the stimulated unit were monitored. If a spike was observed in one of the monitored electrodes within 50 ms of the stimulation pulse, the unit was considered to have responded to the stimulation. The stimulation probability was then calculated as the fraction of times a spike was observed after a stimulation (n = 20).

##### Immunocytochemistry

Cells plated on coverslips were rinsed with PBS before fixation with 4% paraformaldehyde (PFA) for 15 minutes at room temperature. Cells plated on MEA chips were rinsed with PBS three times before PFA fixation (25 minutes, room temperature). All samples were then rinsed with PBS again and blocked for 1 hour at room temperature in PBS with 5% donkey serum and 0.3% Triton X-100 (PBST). The samples were then incubated with primary antibodies (dilutions shown in Table 1) in PBST plus 5% donkey serum overnight at 4 °C. The antibodies were rinsed off the following day with 3 PBS washes and incubated for 1 hour at room temperature with secondary antibodies diluted 1:500 with PBST containing 5% donkey serum. After three additional 5 min PBS rinses, samples were incubated with DAPI (4′,6-diamidino-2-phenylindole) at 1 mg/mL for 10 minutes at room temperature. Samples on coverslips were rinsed two more times in PBS, then mounted using Hydromount (National Diagnostics, HS-106) before imaging. MEA chips were imaged by covering the surface with 10µL PBS and mounting under a glass coverslip.

**Table 1:** Antibody.

| Antibody | Manufacturer | Catalog # | RRID # | Dilution |
| --- | --- | --- | --- | --- |
| MAP2 | Abcam<br>(Boston, MA,<br>USA) | ab5392 | AB_213815<br>3 | 1:2500 |
| APP | Abcam | ab126732 | AB_111317<br>27 | 1:400 |
| AnkG | Millipore<br>(Burlington<br>MA, USA) | MABN66 | AB_274980<br>6 | 1:400 |
| 6E10 | Biolegend (San<br>Diego, CA,<br>USA) | Sig-3920 | AB_256465<br>3 | 1:1000 |
| Synapsin I | Millipore | AB1543 | AB_90757 | 1:500 |

##### Imaging

The coverslips were imaged on a Nikon Eclipse Ti2 Confocal microscope with a 60X (1.4 Numerical Aperture) oil lens and A1R (Andor, 1024µm x 1024µm) camera, X-Cite 120 LED illumination system, and DAPI (400-418nm), FITC 9450-490nm), TRITC (554-570nm) and Cy5 (640-670nm) filter sets controlled by NIS-Elements AR software (5.21.03 64-bit). The immunolabeling on the MEA chips were conducted on a Nikon Eclipse Ni-U inflorescent microscope with a 20X objective and equipped with DAPI (400-418nm), FITC 9450-490nm), TRITC (554-570nm) and Cy5 (640-670nm) filter sets controlled by NIS-Elements AR software [Version 5.02.00 (Build 1266), LO, 64-bit].

### Statistical analysis

To analyze the effect of conditions and time on the cultures compared to the wild type ones, we performed 2-way ANOVA with Tukey correction for multiple tests. If a statistically significant difference was observed in the ANOVA then post-hoc tests were performed to test the difference on specific days. These were carried out as non-parametric two-tailed Mann-Whitney t-tests with Holm-Sidak method to adjust the p-values. All p-values reported in the manuscript are adjusted in this way.

## Acknowledgments

The authors gratefully acknowledge Drs. Michael S. Gold, Ron Hart, and Zhiping Pang for helpful suggestions during the writing of the manuscript. HA thanks Dr. Andrew Schwartz for financial support during the execution of the work and the preparation of the manuscript. The work itself relied on financial support from The University of Pittsburgh Momentum Fund and start-up funds from the University of Pittsburgh, School of Medicine, Dept. of Neurobiology to KH

## SUPPLEMENTAL INFORMATION

### Document S1

**Supp Figure 1**. Representative heatmaps of the firing rate (top), spike amplitude (middle) and active electrodes (bottom) of a single wild type culture recorded every four days from DIV10 to 30.

**Supp Figure 2**. Average number of axonal branches detected in different conditions as a function of DIV. n=10 for WT, n=5 for Aβ, n=8 for lentiAPP, n=6 for R1.40.

**Supp Figure 3**. Conduction speed of action potential propagation in different cultures. Black: wildtype culture (n=10), purple: R1.40 culture (n=6), green: lentiAPP (n=8); red: wildtype cultures treated with 1µM Aβ oligomers (n=5) at different DIVs.

### Document S2

**Supp Movie 1**. Example of a 100 ms recording demonstrating the propagation of an action potential from its initiation site. The same site from a wild type culture was analyzed at DIV 14 and again at DIV 18. The propagation shows that the main axons remain the same but have developed additional branches on DIV 18 compared to DIV 14.

