## Supplemental Figure 1, 2 and 3. for "APP overexpression delays synaptic development and alters neuronal network properties"

**SUPPLEMENTAL INFORMATION**

**Document S1.**

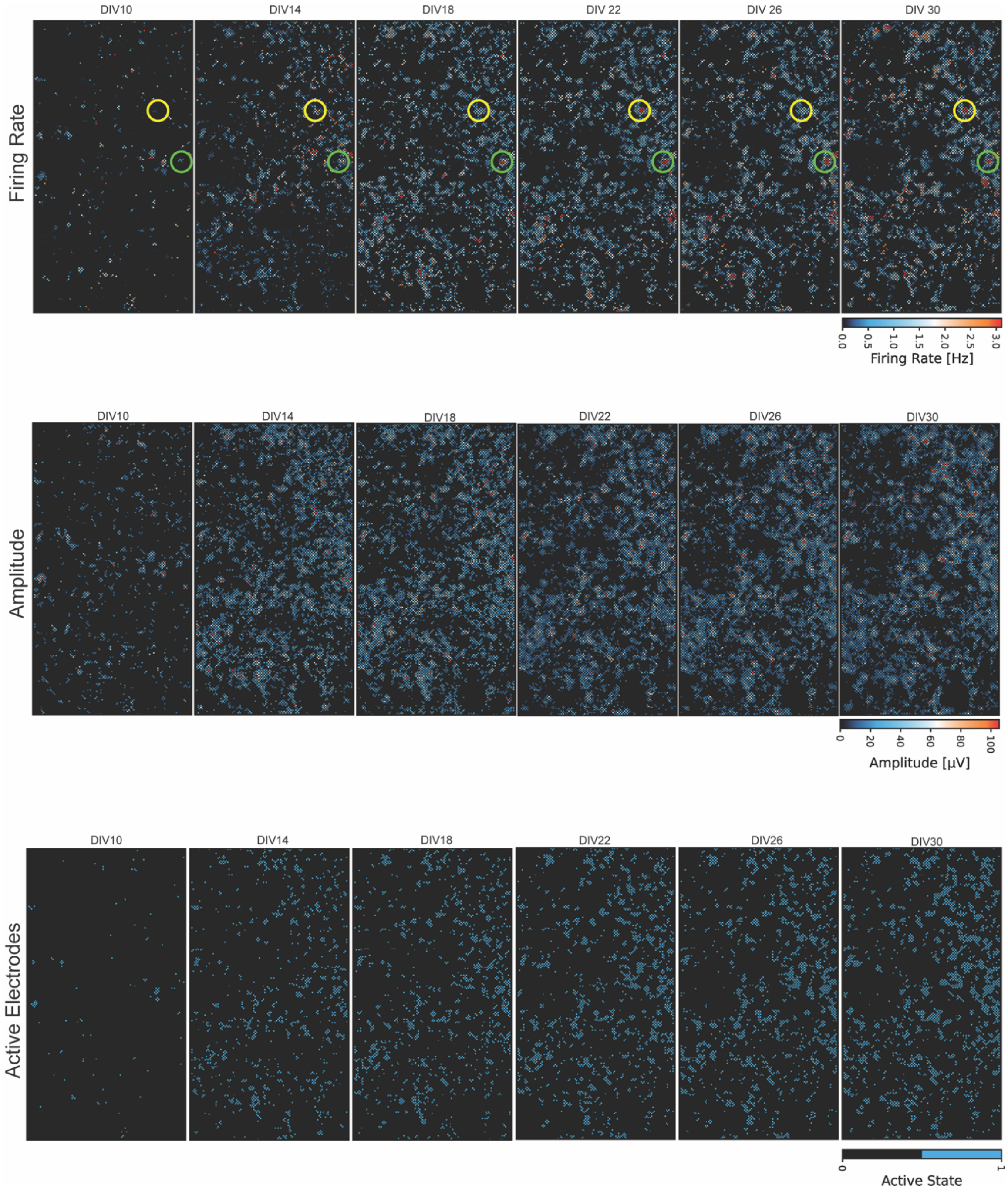


**Supp Figure 1**. Representative heatmaps of the firing rate (top), spike amplitude (middle) and active electrodes (bottom) of a single wild type culture recorded every four days from DIV10 to 30.

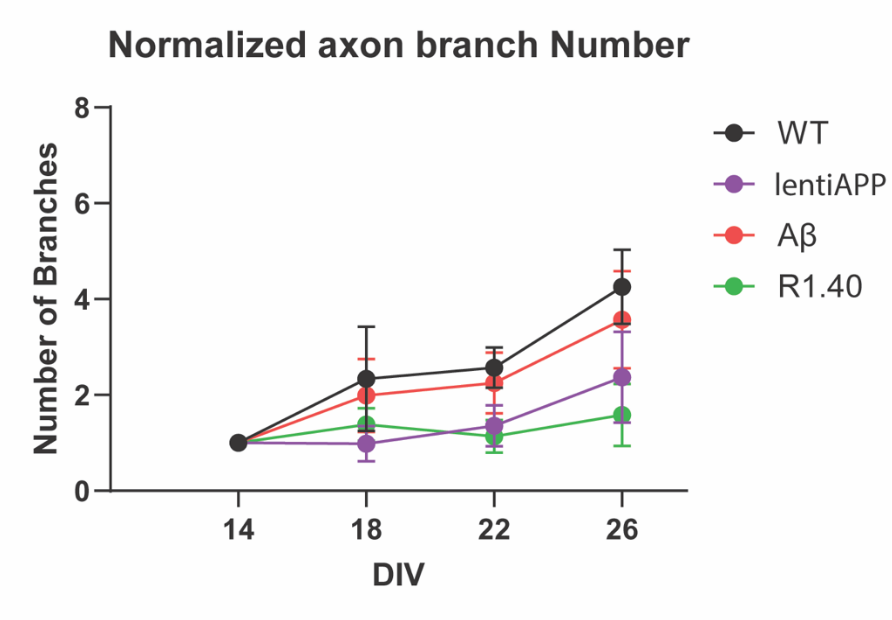


**Supp Figure 2**. Average number of axonal branches detected in different conditions as a function of DIV. n=10 for WT, n=5 for Aβ, n=8 for lentiAPP, n=6 for R1.40.

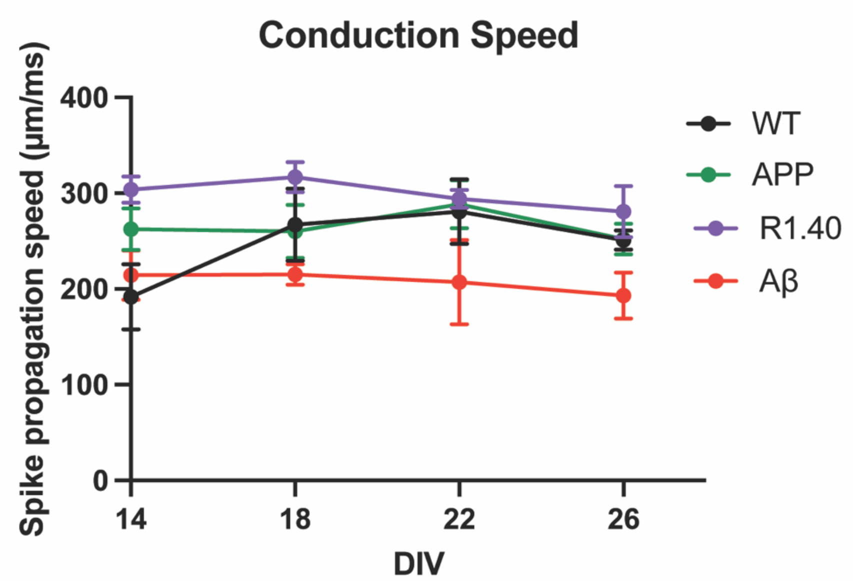


**Supp Figure 3**. Conduction speed of action potential propagation in different cultures. Black: wildtype culture (n=10), purple: R1.40 culture (n=6), green: lentiAPP (n=8); red: wildtype cultures treated with 1µM Aβ oligomers (n=5) at different DIVs.

**Document S2.**

**Supp Movie 1**. Example of a 100 ms recording demonstrating the propagation of an action potential from its initiation site. The same site from a wild type culture was analyzed at DIV 14 and again at DIV 18. The propagation shows that the main axons remain the same but have developed additional branches on DIV 18 compared to DIV 14.
